# Development of Circadian Circuitry Underlying Feeding-Fasting Rhythms in *Drosophila* Larvae

**DOI:** 10.64898/2026.10.23.761381

**Authors:** Nabamita Halder, Christopher McEnaney, Stephanie E. Hood, Amy R. Poe

**Affiliations:** Department of Biological Sciences, University of Arkansas, Fayetteville, AR 72701, USA

## Abstract

Many organisms do not exhibit strong daily behavioral rhythms early in life. The consolidation of sleep and feeding to specific times of the day does not emerge until later in development. Disruptions to circadian behaviors in early life can lead to dysregulation of both neural development and metabolic homeostasis. However, the circuit mechanisms regulating feeding rhythm development are unknown. Here, we characterize the neural circuitry responsible for driving the development of feeding-fasting rhythms in early 3^rd^ instar (L3) *Drosophila* larvae. We find that inactivation of both larval s-LNv circadian pacemaker neurons and *SIFamide* (SIFa) neurons in the *pars intercerebralis* disrupts feeding rhythm development. Additionally, our data indicate feeding rhythms likely emerge due to circuit connections forming between s-LNvs and SIFa neurons at the L3 stage. Finally, we show that reductions in signaling between s-LNv and SIFa neurons interfere with normal behavioral rhythm development in L3. Specifically, RNA interference-induced reductions in *pigment-dispersing factor* (*pdf*) signaling and *SIFa* signaling disrupt diurnal differences in both feeding and sleep, resulting in a loss of behavioral rhythms. Altogether, our findings identify for the first time the clock and clock output circuitry responsible for feeding rhythm regulation in developing organisms.

## Introduction

Molecular circadian clocks are highly conserved across species and begin cycling in early development. However, most organisms early in life have fragmented sleep-wake and feeding-fasting behaviors that are spread out across the circadian day, suggesting that additional maturation steps are required to couple molecular timing to overt behavior (Carmona-Alcocer et al., 2018, Sehgal et al., 1992, Poe et al., 2021, Vallone et al., 2007). For example, newborn infants sleep and feed intermittently throughout the day and night, with consolidated sleep patterns emerging between 3 and 12 months of age (Roffwarg et al., 1966, Davis et al., 2004, Sheldon, 2006). In humans, developmental disruptions to sleep and circadian rhythms frequently co-occur with neurodevelopmental disorders like ADHD and autism (Kotagal, 2015, Hoffmire et al., 2014, Krakowiak et al., 2008). Across species, circadian misalignments in young organisms result in disruptions in adult behaviors, neural development, and physiological functioning (Kotagal, 2015, Doldur-Balli et al., 2022, Cheng et al., 2017).

Understanding how early-life circadian timing interacts with neural development is therefore critical for elucidating the mechanisms shaping lifelong behavioral and physiological function. However, the circuit mechanisms regulating the emergence of behavioral rhythms such as feeding-fasting rhythms remain largely unknown.

Most of our understanding of the fundamental circuit mechanisms regulating feeding-fasting rhythms comes from studies in *Drosophila melanogaster*. Adult flies exhibit diurnal changes in feeding behavior with a peak in feeding in the early daytime and a decrease in feeding at night (Xu et al., 2008). In adult flies, behavioral rhythms including sleep and feeding are regulated through communication between clock neurons and clock output neurons in the *pars intercerebralis* (PI) (Barber et al., 2016, Cong et al., 2015, Martelli et al., 2017, Fulgham et al., 2021). The small ventrolateral neurons (s-LNvs) signal time-of-day information to the rest of the clock network via release of *pigment-dispersing factor* (pdf) (Grima et al., 2004, Stoleru et al., 2004).

Additionally, molecular clock disruptions in the s-LNvs degrade both locomotor activity and feeding rhythms supporting their role in driving behavioral rhythms (Saurabh et al., 2024). Downstream of the clock neurons, clock output neurons in the PI including *insulin-producing cells (IPCs), Diuretic hormone (Dh44)* neurons*, and SIFamide (SIFa)* neurons govern daily behavioral rhythms (Martelli et al., 2017, Barber et al., 2016, Cong et al., 2015, Cavanaugh et al., 2014, King et al., 2017). SIFa neurons release the neuropeptide, *SIFamide*, to regulate feeding rhythms (Martelli et al., 2017, Dreyer et al., 2019, Velazquez et al., 2025) and IPCs release *insulin-like peptide 2* (*dilp2*) to regulate feeding and sleep rhythms (Barber et al., 2016, Cong et al., 2015). While previous studies have characterized the clock neuron-clock output neuron connections that regulate adult behavioral rhythms, few studies have examined how these connections form in the first place. Our previously published work showed that sleep-wake and feeding rhythms emerge at the early 3^rd^ instar (L3) stage (Poe et al., 2023, Poe et al., 2024). L3 larvae exhibit diurnal sleep and feeding patterns similar to adult flies with increased sleep and decreased feeding at night (Poe et al., 2023, Poe et al., 2024).

Additionally, sleep and feeding rhythms in L3 require a molecular clock as they are absent in circadian clock mutants (Poe et al., 2023, Poe et al., 2024). While our published work provides the first evidence that multiple behavioral rhythms emerge at the early L3 stage (Poe et al., 2024), the neurons and signaling pathways responsible for regulating feeding-fasting rhythm development remain unknown.

Here, we examined the role of larval clock and clock output neurons in regulating larval feeding rhythm development. We demonstrate that larval *pdf*-expressing s-LNv neurons regulate the emergence of feeding-fasting rhythms in L3 larvae. Additionally, we determine that larval feeding rhythms are primarily regulated through the actions of SIFa neurons in the PI. Further, we show that a functional connection forms between s-LNvs and SIFa neurons at the early L3, but not prior to this time, to drive feeding rhythms. Finally, we find that disruptions of neuropeptidergic signaling between s-LNv and SIFa neurons affect both sleep-wake and feeding-fasting rhythms in L3 larvae, suggesting that larval behavioral rhythms are regulated by multiple neuromodulatory mechanisms. Together, our data demonstrate that larval behavioral rhythms emerge at the L3 stage through the establishment of new circuit and signaling connections between clock and clock output neurons. These findings provide insight into how early-life circadian circuits and behavioral rhythms are established during development.

## Results

### Inactivation of pdf-expressing s-LNvs and SIFamide neurons disrupts feeding rhythms

Feeding-fasting rhythms emerge at the early 3^rd^ instar (L3) stage in *Drosophila* (Poe et al., 2024). To determine if feeding-fasting rhythm development is regulated by larval s-LNv neurons, we inhibited *pigment-dispersing factor*-expressing (Pdf) s-LNv neurons by expressing the inwardly rectifying potassium channel *Kir2.1* (Baines et al., 2001). Using an established blue dye-based feeding assay (Poe et al., 2024), we then measured food intake in the subjective day (Circadian Time [CT] 7) and the subjective night (CT19) in developmentally age-matched early L3 larvae (72 hours after egg laying [AEL]). We found that silencing the *pdf*-expressing neurons caused a decrease in daytime feeding amount and an increase in nighttime feeding amount resulting in a loss of feeding rhythms (Figure 1A). However, genetic control larvae showed the expected increase in food intake at CT7 compared to CT19 (Figure 1A). We next examined whether Pdf(+) inhibition also affects sleep-wake rhythm development in early L3. Kir2.1 expression in the Pdf(+) neurons did not affect diurnal rhythms in sleep with larvae showing the expected increase in sleep amount at night but did cause a slight decrease in sleep bout length at CT19 compared to genetic controls (Figures 1B-D). Together, these data suggest that *pdf*-expressing s-LNvs neurons play a minor role in regulating larval sleep architecture but are primarily responsible for regulating feeding rhythm development.

**Figure 1.**
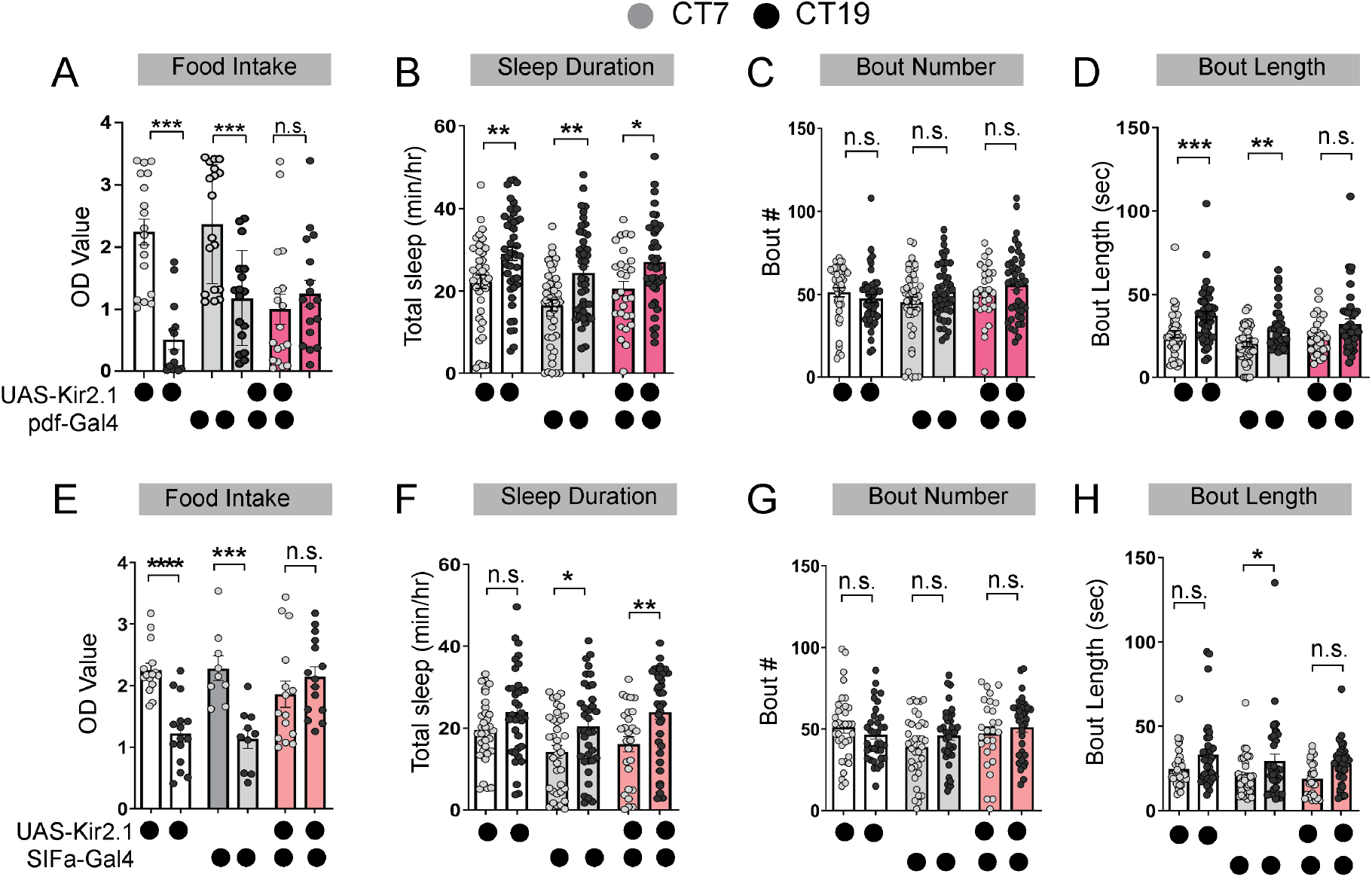
Inactivation of s-LNv and SIFa neurons disrupts feeding rhythms. **(A)** Feeding amount in L3 expressing *pdf*-Gal4>*UAS-Kir2.1* and genetic controls at CT7 and CT19. **(B-D)** Sleep duration (B), bout number (C) and bout length (D) of L3 expressing *pdf*-Gal4>*UAS-Kir2.1* and genetic controls at CT7 and CT19. **(E)** Feeding amount in L3 expressing *SIFa-*Gal4>*UAS-Kir2.1* and genetic controls at CT7 and CT19. **(F-H)** Sleep duration (F), bout number (G), and bout length (H) in L3 expressing *SIFa-*Gal4>*UAS-Kir2.1* and genetic controls at CT7 and CT19. The subjective day (CT7) is shown in light gray circles and the subjective night (CT19) is shown in black circles. (A, E), n = 20 replicates (13 larvae/replicate); (B-D), (F-H), n = 40-45 larvae. For this and all other figures unless otherwise specified, data are represented as mean ± SEM; *p<0.05, **p<0.01, ***p<0.001, ****p<0.0001, n.s.= not significant. (A-H) One-way ANOVA followed by Tukey’s multiple comparisons test.

Next, to determine if *SIFamide* (SIFa) neurons in the larval PI regulate feeding rhythms in L3 larvae, we expressed Kir2.1 in SIFa neurons using *SIFa*-Gal4 and examined food intake at CT7 and CT19. Kir2.1 expression in SIFa neurons caused an increase in the nighttime feeding amount, but no effect on daytime feeding, resulting in a loss of diurnal differences in feeding (Figure 1E). In contrast, the genetic controls showed the expected increase in feeding amount at CT7 compared to CT19 (Figure 1E). Finally, we examined whether inhibition of SIFa neurons affects sleep-wake rhythms. We found that silencing *SIFa*-expressing neurons did not disrupt sleep-wake rhythm development with SIFa inhibition larvae showing a significant increase in sleep amount at CT19 compared to CT7 and mild increases in nighttime sleep bout duration (Figures 1F-H). Altogether, these data suggest that SIFa neurons primarily regulate larval feeding-fasting rhythms.

### s-LNv neurons functionally connect with SIFa neurons in early L3

In adult flies, s-LNv and SIFa neurons form circuit connections to regulate rhythmic behaviors (Reinhard et al., 2024). We next asked whether these two neurons form circuit connections at the early L3 stage. To test whether these neurons are functionally connected in early L3, we expressed adenosine 5’-triphosphate (ATP)-gated P2X2 receptors in *pdf*-expressing neurons and *UAS-GCaMP7f* in SIFa neurons. When pdf(+) neurons were activated with ATP application to the dissected brains, we observed an increase in calcium in SIFa neurons in early L3 (Figures 2A and B). However, surprisingly, we did not observe an increase in calcium in SIFa neurons in 2^nd^ instar (L2) larvae (Figures 2C and D) indicating that functional connections form between s-LNv and SIFa neurons at the L3 stage, but not prior to this time. Given that the functional connection forms at the same time as feeding rhythms (Poe et al., 2024), our data suggest that the s-LNv-SIFa neuron circuit plays a role in the emergence of feeding rhythms development in *Drosophila* larvae.

**Figure 2.**
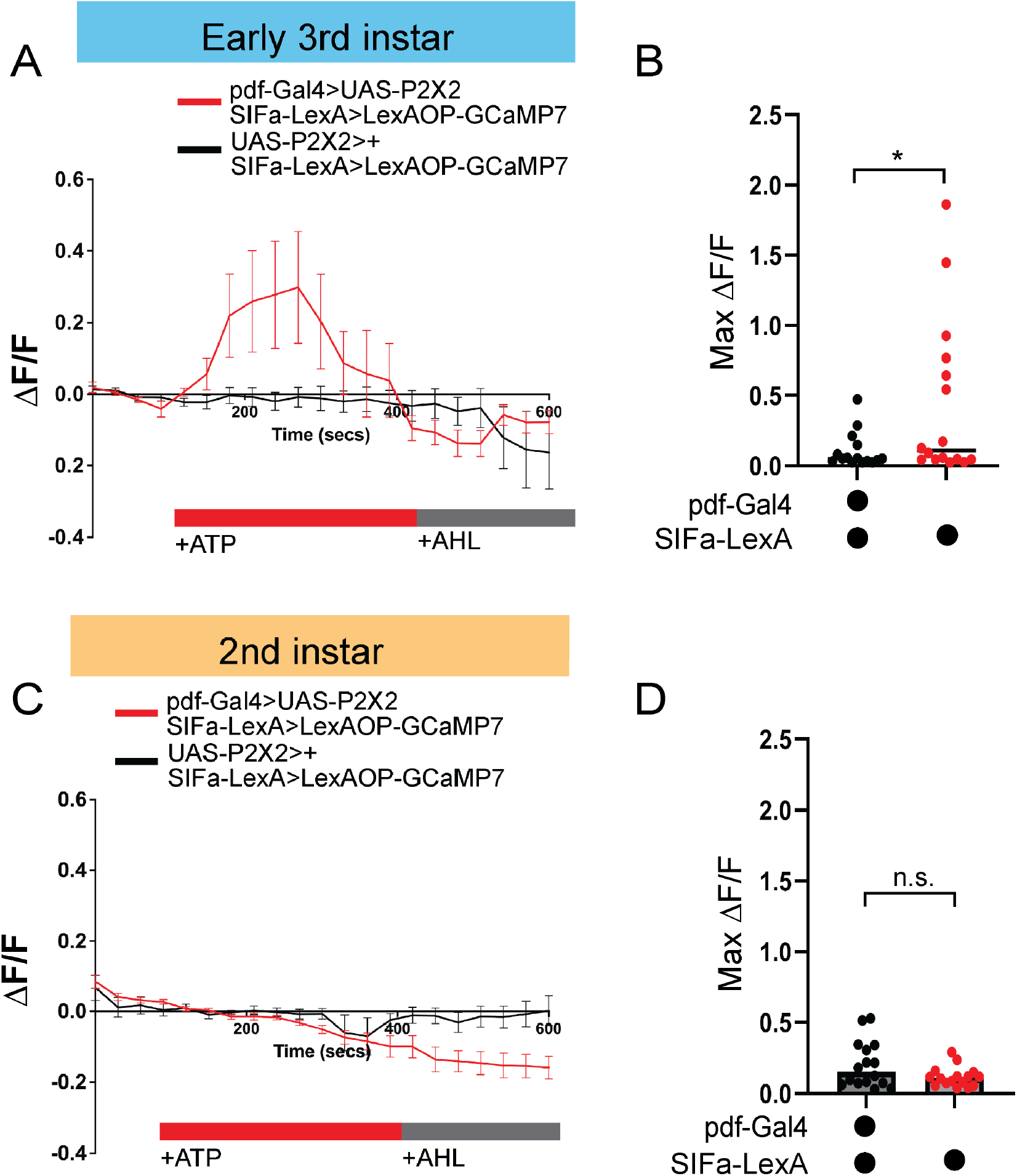
s-LNv neurons functionally connect with SIFa neurons in early L3. **(A)** GCaMP7 signal in SIFa neurons with activation of s-LNv neurons in L3. The red bar indicates adenosine 5′-triphosphate (ATP) application and the gray bar indicates artificial hemolymph (AHL) application. **(B)** Maximum GCaMP7 change (ΔF/F) for individual cells in L3. **(C)** GCaMP7 signal in SIFa neurons with activation of s-LNv neurons in L2. The red bar indicates ATP application and the grey bar indicates AHL application. **(D)** Maximum GCaMP7 change (ΔF/F) for individual cells in L2. A-D, n = 10–15 cells; 10 brains. (B, D) Unpaired two-tailed Student’s t-test.

### Pdf and SIFamide signaling between s-LNv and SIFa neurons is required for feeding rhythms

Thus far, our data suggest that inactivation of both s-LNv neurons and SIFa neurons disrupt feeding rhythms at the L3 stage. We next sought to determine the molecular signals used in the s-LNv-SIFa circuit to drive feeding rhythms. To first determine if *Pdf* release from s-LNvs is required, we used RNA interference (RNAi) to knockdown *pdf* expression using *pdf*-Gal4. We found that *pdf* knockdown disrupted feeding rhythms due to reductions in daytime (CT7) feeding amount compared to genetic controls (Figure 3A). In contrast to our *pdf* inhibition experiments, however, we observed a loss of sleep-wake rhythms in *pdf* knockdown larvae with a decrease in nighttime sleep amount, bout number and bout length (Figures 3B-D). Next, we used RNAi to knockdown *SIFa* expression using *SIFa*-Gal4 and observed a loss of feeding rhythms driven by an increase in nighttime (CT19) feeding amount compared to genetic controls (Figure 3E). We also observed a loss of sleep-wake rhythms in *SIFa* knockdown larvae due to an increase in both daytime sleep duration and the number of daytime sleep episodes (bouts) (Figures 3F-H). Altogether, these data show that knockdown of either *pdf* or *SIFa* expression disrupts both larval feeding and sleep-wake rhythms, suggesting that release of these neuropeptides likely affects both sleep-regulatory neurons and feeding-regulatory neurons.

**Figure 3.**
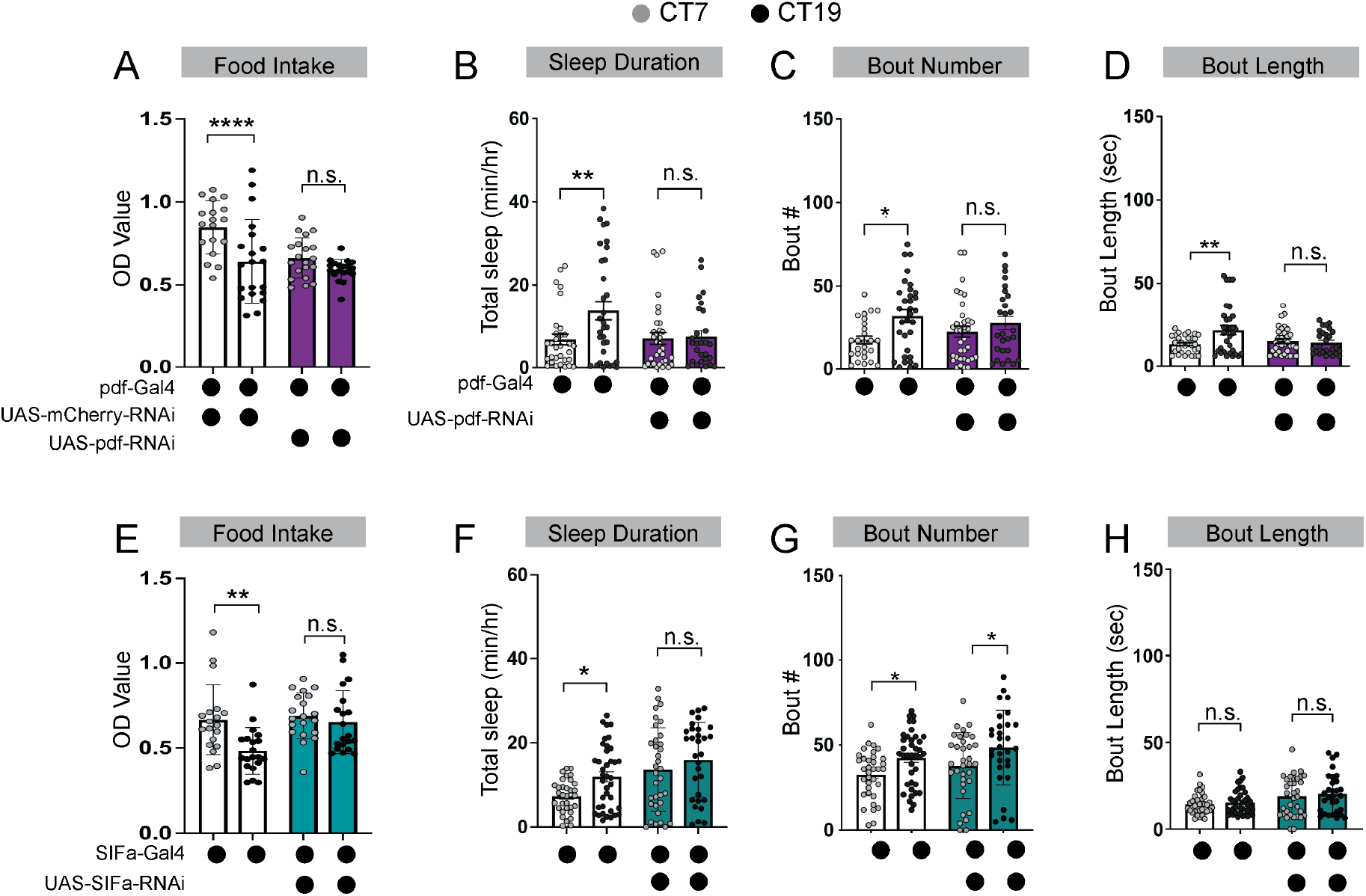
pdf and SIFa release are required for feeding and sleep-wake rhythms. **(A)** Feeding amount in L3 expressing *pdf*-Gal4>*UAS-pdf-RNAi* and genetic controls at CT7 and CT19. **(B–D)** Sleep duration (B), bout number (C), and bout length (D) of L3 expressing *pdf*-Gal4>*UAS-pdf-RNAi* and genetic controls at CT7 and CT19. **(E)** Feeding amount in L3 expressing *SIFa-*Gal4*>UAS-SIFa-RNAi* and genetic controls at CT7 and CT19. **(F–H)** Sleep duration (F), bout number (G), and bout length (H) of L3 expressing *SIFa-*Gal4*>UAS-SIFa-RNAi* and genetic controls at CT7 and CT19. (A, E), n=20 replicates (13 larvae/replicate); (B-D), (F-H), n = 40–45 larvae. (A-H) One-way ANOVA followed by Tukey’s multiple comparisons test.

Finally, to test whether *pdf*-expressing s-LNvs and SIFa neurons communicate with each other to modulate feeding rhythms, we used RNAi to knockdown expression of the *SIFamide receptor* (*SIFaR*) in the s-LNvs. Consistent with our prior *SIFa* knockdown results, we observed a loss of feeding rhythms driven by a decrease in daytime feeding amount in *SIFaR* knockdown larvae (Figure 4A). Additionally, we observed a loss of diurnal differences in sleep with *SIFaR* knockdown larvae showing a reduction in total nighttime sleep duration, bout number, and bout length (Figures 4B-D). Next, we examined whether SIFa neurons require *pdf* signals to drive feeding and sleep rhythms. We knocked down expression of the *pdf receptor* (*pdfR*) in SIFa neurons and examined feeding rhythms. We found that *pdfR* knockdown larvae showed an increase in feeding amount at both CT7 and CT19 leading to a loss of feeding rhythms (Figure 4E). *pdfR* knockdown larvae also exhibited a loss of sleep-wake rhythms driven by a significant increase in daytime sleep duration and trend towards more daytime sleep bouts (Figures 4F-H). Altogether, our findings suggest that a developmentally timed circuit between s-LNv and SIFa neurons use peptidergic signaling to regulate both sleep-wake and feeding rhythm development.

**Figure 4.**
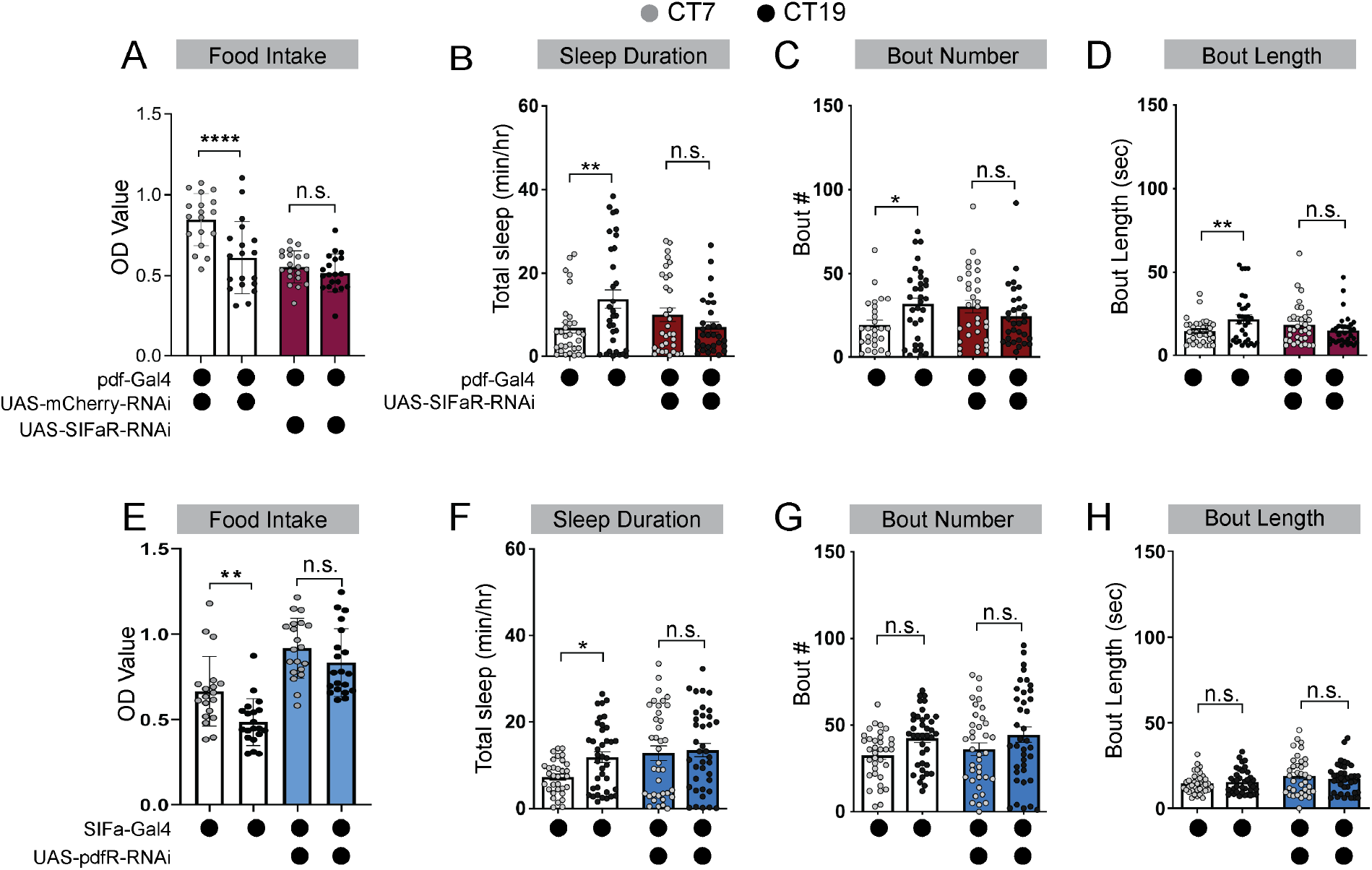
Pdf receptor and SIFamide receptor signaling is required for feeding rhythms in early L3. **(A)** Feeding amount in L3 expressing *pdf-*Gal4*>UAS-SIFaR-RNAi* and genetic controls at CT7 and CT19. **(B–D)** Sleep duration (B), bout number (C), and bout length (D) of L3 expressing *pdf-*Gal4*>UAS-SIFaR-RNAi* and genetic controls at CT7 and CT19. **(E)** Feeding amount in L3 expressing *SIFa-*Gal4*>UAS-pdfR-RNAi* and genetic controls at CT7 and CT19. **(F–H)** Sleep duration (F), bout number (G), and bout length (H) of L3 expressing *SIFa-*Gal4*>UAS-pdfR-RNAi* and genetic controls at CT7 and CT19. (I) L3 brain showing *pdfR*-Gal4>*UAS-mCD8-GFP* expression in PI region of the brain. Arrows points to SIFa cell bodies. (J) L3 brain showing *SIFaR*-Gal4>*UAS-mCD8-GFP* expression in s-LNv cell bodies. (A, E), n = 20 replicates (13 larvae/replicate); (B-D), (F-H), n = 40–45 larvae. (A-H) One-way ANOVA followed by Tukey’s multiple comparisons test.

## Discussion

The proper development of circadian rhythms plays a critical role in nervous system formation. However, the circuit mechanisms regulating behavioral rhythm emergence in developing organisms are not well understood. Here, we identify a circuit between s-LNv *(Pdf+)* clock neurons and the SIFa neurons that plays a primary role in regulating feeding rhythms in *Drosophila* larvae. We show that silencing s-LNv *(Pdf+)* neurons disrupts feeding rhythms while sleep-wake rhythms remain largely intact.

Similarly, silencing SIFa neurons disrupts feeding rhythms, but does not affect sleep-wake rhythms, suggesting that SIFa neurons function as a dedicated circadian output for feeding rhythms. Our data further demonstrate that a functional connection between s-LNv and SIFa neurons forms at the early L3 stage but is absent in L2 larvae, suggesting that the emergence of feeding rhythms is under tight developmental control. Finally, our RNAi experiments suggest that signaling between s-LNvs and SIFa neurons is likely necessary for both sleep-wake and feeding rhythm maturation. RNAi knockdown of either *pdf* or *SIFaR* in the s-LNvs decreases daytime feeding and nighttime sleep amounts resulting in a loss of both rhythms. Additionally, knockdown of either *SIFa* or *pdfR* in SIFa neurons increases both daytime sleep and nighttime feeding resulting in a loss of behavioral rhythms. These findings suggest that the release of neuropeptides from s-LNv and SIFa neurons is necessary for behavioral rhythm development in L3 larvae.

Together, with our previously published work (Poe et al., 2023, Poe et al., 2024), our findings support a model in which the formation of circuit connections between clock neurons and clock output neurons gates the onset of circadian behaviors in developing *Drosophila* larvae. Interestingly, our s-LNv/SIFa inhibition and RNAi knockdown experiments suggest that s-LNv and SIFa neurons might regulate distinct aspects of feeding and sleep timing as we observe time-of-day effects on these behaviors in those manipulations. As the s-LNvs act as the morning cells to drive anticipatory activity in adult flies (Grima et al., 2004, Stoleru et al., 2004), our data suggest that these neurons serve developmentally conserved roles in regulating daytime behaviors in larvae. In contrast, our data suggest that SIFa neurons may predominately regulate evening feeding behaviors. These findings raise the intriguing possibility that the larval brain utilizes a complex network of peptidergic signals between clock neurons and clock outputs to precisely consolidate sleep, waking, feeding, and fasting to certain times of day. In support of this, our previous work has shown that sleep-wake rhythm development is regulated by the formation of a circuit between DN1a and Dh44 neurons and *CCHamide-1* signaling between these neuronal populations (Poe et al., 2023).

However, it is not known if these neurons also receive signals from the s-LNv and SIFa neurons to regulate sleep and arousal. Future work will determine whether the DN1a and Dh44 neurons receive other peptidergic signals like *pdf* or *SIFa* to modulate sleep-wake rhythm development. Nevertheless, our data provide the first insights into how these signals modulate the development of multiple behavioral rhythms.

Previous studies in adult *Drosophila* demonstrate that s-LNv and SIFa neurons play a role in regulating feeding-fasting rhythms (Martelli et al., 2017, Dreyer et al., 2019, Velazquez et al., 2025). In adult flies, s-LNv neurons regulate circadian behaviors through the release of *pdf* (Grima et al., 2004, Stoleru et al., 2004), and disruption in the central clock neurons disrupts both locomotor and feeding rhythms (Saurabh et al., 2024). Downstream of the clock, SIFa neurons in the PI serve as a dedicated output center for feeding-fasting rhythms, with SIFamide and its receptor required for rhythmic food intake (Martelli et al., 2017, Dreyer et al., 2019, Velazquez et al., 2025). Our findings expand on these prior studies by providing new insights into how s-LNv-SIFa connections first develop.

In adults, feeding rhythms are also regulated by a diverse array of cells in both the brain and peripheral tissues. For example, insulin-producing cells (IPCs) in the PI release *insulin-like peptide 2* (*dilp2*) to regulate both feeding and sleep rhythms (Barber et al., 2016, Cong et al., 2015). s-LNvs functionally modulate IPCs in a time-of-day manner (Hameed et al., 2026), providing further support for their role in feeding behavior regulation. Recent studies also demonstrate that SIFa, Dh44, & IPCs likely form synaptic connections with each other (McKim et al., 2026) raising the possibility that communication between each neuronal population in the PI may modulate larval behavior rhythms. Future studies will examine whether larval IPCs also regulate sleep-wake and feeding rhythms through the central clock or other clock output neurons.

Finally, adult fly feeding rhythms are also regulated by molecular clocks in other peripheral tissues such as the gut and fat body (Xu et al., 2008, Fulgham et al., 2021, Wu and Ja, 2026). While recent studies suggest that rhythmic expression of clock genes does not begin in the intestine until the adult stage (Parasram et al., 2024), the development of the peripheral clock is an understudied process. Future work will examine whether disruptions of peripheral clock functioning in the larval fat body also disrupt larval feeding rhythms. These studies will open new lines of investigation into the brain-body mechanisms regulating larval feeding rhythm development.

Broadly, our findings imply that the emergence of behavioral rhythms is driven by developmental changes in circuit connections and communication between clock neurons and clock output neurons. In mammals, feeding rhythms emerge gradually with rodents displaying adult feeding patterns by 30-35 days after birth (de Castro and Balagura, 1975) and human infants showing consolidated feeding periods in the first few months of life (Roffwarg et al., 1966, Davis et al., 2004). These rhythms largely develop due to the establishment of neural projections between the mammalian suprachiasmatic nucleus (SCN) and neurons in the hypothalamus (Bechtold and Loudon, 2013). The precise development of these circuits is essential, as disruptions in early-life circadian rhythms are a common comorbidity in human neurodevelopmental and metabolic disorders, including ADHD, obesity, and type 2 diabetes (Kotagal, 2015, Hoffmire et al., 2014, Mariano et al., 2023, Valadas et al., 2018, Cirelli, 2006).

Therefore, our results provide critical insight into the conserved mechanisms in which developing organisms consolidate their feeding to certain times of day to achieve proper metabolic homeostasis. In line with this, our prior work suggests that changes in metabolic needs drive sleep-wake rhythm development (Poe et al., 2024) indicating that the consumption of nutrients to certain times of day is a critical driver of behavioral rhythm development. Future studies will examine the effects of improper nutritional environments on feeding-sleep wake rhythm crosstalk.

## Acknowledgements

We thank Chris Huang, Madeline Lee, and Houston Collard for their assistance with larval sample collections. We thank members of the Poe lab for helpful discussions and input.

## Author contributions

Conceptualization, N.H., C.M., A.R.P.; Investigation, N.H., C.M., S.E.H., A.R.P.; Writing – Original Draft, N.H., C.M., and A.R.P.; Writing – Review & Editing, all authors; Project supervision and funding, A.R.P.

## Data and Materials Availability

All data needed to evaluate the conclusions in the paper are present in the paper.

## Materials and Methods

### Fly Stocks

The following fly lines used were maintained as lab stocks: iso31, UAS-Kir2.1 (Baines et al., 2001), SIFa-Gal4 (Terhzaz et al., 2007), pdf-Gal4 (Renn et al., 1999), and LexAOP-GCamP7f UAS-P2X2 (Yao et al., 2012). The following lines were obtained from the Bloomington Drosophila Stock Center: UAS-mCherry RNAi (35785), UAS-SIFa RNAi (60484), UAS-SIFaR RNAi (44068), UAS-Pdf RNAi (25802), and UAS-Pdfr RNAi (42508).

### Larval rearing

Larval rearing was performed as described previously (Poe et al., 2023, Poe et al., 2024). Briefly, adult flies were maintained at 25°C on standard molasses-based food in a 12:12-hour light: dark (LD) cycle. For experiments performed in the subjective evening (CT12 or CT18), adult flies were maintained on a 12:12 reverse (DL) cycle.

Synchronized early third instar larvae (L3) were collected by placing adult flies in an embryo collection cage (Genesee Cat# 59-100). The eggs were then collected on petri dishes containing molasses-based food with yeast paste on top. The larvae were allowed to develop at 25°C on these plates for 3 days and kept at 25°C for the duration of the experiment.

### Larval feeding assays

Larval food intake was measured using a blue dye-based feeding assay adapted and modified from (Poe et al., 2024). For the assay, early 3^rd^ instar larvae were collected and starved for 1 hour in petri-dishes with a moist Kimwipe at 25°C. Following starvation, 13 larvae per sample (n=20 replicates) were placed in a petri-dish containing blue-dyed agar 2% (w/v), sucrose 1.2% (w/v), apple juice 25% (v/v) and blue-dye 2% (v/v) with blue yeast paste on top for 4 hours at 25°C in constant darkness. The blue yeast paste was prepared using active dry yeast (Genesee, cat. #62-103), water and blue food dye (Carolina Biological Supply, cat. #898032) in the ratio 5:9:1 respectively. After the 4-hour feeding period the larvae are washed in water to remove any residual blue yeast paste and then transferred into 1.5ml microcentrifuge tubes and stored at - 80°C freezer. The larvae were homogenized using a Kimble Kontes Pellet Pestle Cordless Motor (Fisher, cat. # K749540-0000) in 300 µl cold ddH_2_O and centrifuged at 13000 rpm for 5 mins at 4°C. The supernatant was analyzed at OD629 using an Agilent BioTek microplate reader, and food intake was quantified as the OD value of each sample.

### Larval sleep assays

Larval sleep experiments were performed as described previously (Poe et al., 2023, Poe et al., 2024). Molting 3^rd^ instar larvae were placed into individual wells of the LarvaLodge containing 95 µl of 3% agar and 2% sucrose media covered with a thin layer of yeast paste. The LarvaLodge was covered with a transparent glass sheet (Fayetteville Glass Co., 5 × 4 × 1/8 in; length × width × thickness) and placed into a Percival dual-chamber incubator at 25°C for imaging. Experiments were performed in the dark.

### LarvaLodge image acquisition and processing

Image acquisition occurred every 6 seconds using an Imaging Source DMK 23UP031 camera (The Imaging Source, 2592 x 1944 pixels) equipped with a Fujinon lens (HF12.5SA-1, 1:1.4/12.5 mm, Fujifilm Corp., Japan) and a Hoya 49mm R72 IR filter. To acquire time-lapse images, we used IC Capture 4 (The Imaging Source) software.

Images were analyzed using a custom-written MATLAB script as described in (Churgin et al., 2019, Szuperak et al., 2018). Temporally adjacent images were subtracted to generate maps of pixel value intensity change. A binary threshold was set such that individual pixel intensity changes that fell below 45 gray-scale units within each well were set equal to zero (“no change”) to eliminate noise. Pixel changes greater than or equal to threshold value were set equal to one (“change”). Activity was then calculated by taking the sum of all pixels changed between images. Sleep was defined as an activity value of zero between frames. For sleep experiments performed at certain circadian times, total sleep in the 2^nd^ hour after the molt to third instar was summed.

### P2X2 imaging

P2X2 mediated calcium imaging was performed as previously described in (Poe et al., 2023). Larval brains were dissected in artificial hemolymph (AHL) buffer containing (in mM): 108 NaCl, 5 KCl, 2 CaCl_2,_ 8.2 MgCl_2,_ 4 NaHCO3, 1 NaH2PO4.H2O and 5 Trehalose, 10 Sucrose and 5 HEPES (pH= 7.5). Dissected brains were mounted on a 12-mm circular glass coverslip (Carolina Cover Glasses, cat. #633029) within a horizontal perfusion chamber (Automate Scientific, cat. #SD-PC-H) and continuously perfused with AHL using a gravity-fed ValveLink pinch valve perfusion system (Automate Scientific, cat. # 17-pp-24).

Baseline neural activity (GCaMP7) was recorded for 1min with AHL perfusion, then for 2 min with perfusion of 2.5 mM ATP in AHL (pH=7.5), and then for 2 min with perfusion of AHL. Time-lapse imaging was performed on a ZEISS LSM 900 confocal microscope using a 20x inverted objective at 512 x 512 pixel resolution with twelve-bit image depth. Z-stacks were acquired every 5 sec for 5 min. Fluorescence intensity and image processing was performed in ImageJ as previously described (Poe et al., 2023). Briefly, a max intensity Z projection was applied to each time series, and fluorescence intensity was measured from manually selected ROI of individual neurons in ImageJ. The fluorescence traces over time were normalized using the equation: ***Δ***F/F = (F_n_ − F₀)/F₀, where F_n_= fluorescence intensity at time point n and F₀ is the average fluorescence value during the 1 min baseline recording. All the analyses were conducted blind to experimental conditions.

### Statistical analysis

All statistical analyses were performed using GraphPad Prism software. Comparison between two groups were measured using two-tailed unpaired t-tests. For datasets involving more than two groups, ordinary one-way ANOVA was applied, followed by Tukey’s multiple comparison tests. For comparisons between different groups in the same analysis, ordinary one-way ANOVAs followed by Sidak’s multiple comparisons tests were used. For comparisons between time and genotype, two-way ANOVAs followed by Sidak’s multiple comparisons tests were used. Statistical significance was defined as *p<0.05, **p<0.01, and ***p<0.001. For live imaging calcium activity was measured from ∼10-15 neurons across 10 brains per condition.

